# Regulatory divergence as a mechanism for X-autosome incompatibilities in *Caenorhabditis* nematodes

**DOI:** 10.1101/2022.06.28.498049

**Authors:** Athmaja Viswanath, Asher D. Cutter

## Abstract

The world’s astounding biodiversity results from speciation, the process of formation of distinct species. Hybrids between species often have reduced fitness due to negative epistatic interactions between divergent genetic factors, as each lineage accumulated substitutions independently in their evolutionary history. Such negative genetic interactions can manifest as gene misexpression due to divergence in gene regulatory controls from mutations in *cis*-regulatory elements and *trans*-acting factors. Gene misexpression due to differences in regulatory controls can ultimately contribute to incompatibility within hybrids through developmental defects such as sterility and inviability. We sought to quantify the contributions of regulatory divergence to post-zygotic reproductive isolation using sterile interspecies hybrids of two *Caenorhabditis* nematodes: *C. briggsae* and *C. nigoni*. We analysed previous transcriptome profiles for two introgression lines with distinct homozygous X-linked fragments from *C. briggsae* in a *C. nigoni* genomic background that confers male sterility, owing to defects in spermatogenesis (Li et al. 2016). Our analysis identified hundreds of genes that show distinct classes of non-additive expression inheritance and regulatory divergence. We find that these non-overlapping introgressions affect many of the same genes in the same way and demonstrate that the preponderance of transgressive gene expression is due to regulatory divergence involving compensatory and joint effects of *cis-* and *trans*-acting factors. The similar transcriptomic responses to non-overlapping genetic perturbations of the X-chromosome implicate multiway incompatibilities as an important feature contributing to hybrid male sterility in this system.

**Significance statement:** The genetic causes of intrinsic post-zygotic reproductive isolation can arise from hybrids experiencing negative gene regulatory interactions. In *Caenorhabditis* nematodes, hybrid male sterility involves X-autosome incompatibilities that affect small-RNA regulatory pathways. We sought to understand the role of gene regulatory divergence as a related contributor to hybrid misexpression by analyzing transcriptomes of sterile males from two hybrid introgression lines, each containing distinct X-linked fragments from *C. briggsae* in a *C. nigoni* genomic background. We show that gene misexpression occurs due to extensive joint divergence of *cis-* and *trans*-acting regulatory factors and provide evidence for multiway incompatibilities as an important feature of this system.

## INTRODUCTION

Speciation is the process of formation and maintenance of distinct species through the evolution of reproductive isolation. Events that split populations into separated groups, with restricted gene flow between those groups, can eventually lead to evolution of genetically intrinsic reproductive isolation capable of maintaining these groups as distinct species (Coyne & Orr 2004). Intrinsic postzygotic barriers take the form of developmental defects like hybrid sterility or hybrid inviability that can result from dysfunctional genotype-genotype interactions (Coyne & Orr 1998; Wolf et al. 2010; Coughlan & Matute 2020; Cutter & Bundus 2020). Multiple genetic mechanisms can create such incompatibility, including chromosomal rearrangements, gene loss and duplication, repetitive transposable element composition and activity, sequence pairing problems due to divergence, and negative epistatic interactions between two or more loci (i.e., Dobzhansky-Muller incompatibilities, DMIs) (Orr 1995; Presgraves 2010; Maheshwari & Barbash 2011). Such hybrid incompatibilities can involve distinct genomic compartments, including cyto-nuclear incompatibilities or X-autosome incompatibilities (Turelli & Moyle 2007; Bundus et al. 2018). DMIs also may be genetically simple or require more complex interactions of three or more genetic factors, with such multi-way incompatibilities expected to accumulate more rapidly over time (Orr 1995; Kondrashov 2003; Satokangas et al. 2020). Incompatibilities also can arise from protein-protein interactions or from interactions involving gene regulatory regions that yield misexpression in hybrids (Mack & Nachman 2017). The importance of regulatory divergence as an underlying mechanism of hybrid incompatibility, however, remains to be fully elucidated, especially with respect to sex-biases in hybrid dysfunction (Mack & Nachman 2017).

The heterogametic sex, males in *Caenorhabditis* nematodes and many other taxa, is disproportionately afflicted by F1 hybrid sterility and inviability (Delph & Demuth 2016; Cutter 2018). Despite the prevalence of this phenomenon, termed Haldane’s rule (Haldane 1922), there is limited consensus for a leading cause among the several hypotheses proposed to explain it (including dominance theory, faster X theory, faster male theory, and meiotic drive (Wolf et al. 2010; Delph & Demuth 2016). To the extent that the developmental origins of Haldane’s rule involves gene misexpression, regulatory divergence may play a crucial role in the manifestation of sex-biased hybrid incompatibility (Mack & Nachman 2017). Moreover, a wide range of taxa show significant misexpression in interspecies hybrids of male-biased genes (Michalak 2003; Ranz et al. 2004) and spermatogenesis-related genes (Ferguson et al. 2013; Sundararajan & Civetta 2011), and disproportionate misexpression of X-linked genes in sterile hybrids (Lu et al. 2010; Oka & Shiroishi 2014; Gomes & Civetta 2015). These patterns of gene expression suggest that regulatory divergence of sex-biased genes and X-linked genes are instrumental in producing the dysfunction of developmental programs that lead to hybrid sterility (Coyne & Orr 2004; Llopart 2012; Coolon et al. 2015).

Misexpression of a given gene in hybrids can arise due to divergence in regulatory controls caused by mutations in *cis*-regulatory elements encoded close to the gene and by changes in *trans*-acting factors that are encoded elsewhere in the genome (Landry et al. 2007; Mack & Nachman 2017). In F1 hybrids, in which all alleles experience a common *trans*-acting environment, any difference in expression between the two alternate alleles of a gene (allele specific expression) indicates that *cis*-regulatory differences exist between the parents at that locus. If, instead, the gene’s expression differs in F1 hybrids compared to the parents in the absence of F1 allele-specific expression differences, then divergence of *trans*-acting factors must be the cause (Wittkopp et al. 2004). Commonly, however, studies of hybrids find that *cis*-acting changes are compensated by changes in *trans*-acting factors despite similar overall gene expression between species (Takahasi et al. 2011; Goncalves et al. 2012; Mack & Nachman 2017). Compensatory *cis-trans* regulatory evolution like this represents one way that molecular evolution can accrue despite stabilizing selection generally being pervasive to conserve overall gene expression levels between species (Gilad et al. 2006; Signor & Nuzhdin 2018). This approach of assessing overall and allele-specific expression with hybrid organisms has permitted quantification of genome-wide *cis-* and *trans-*acting regulatory divergence in plants, fungi, insects, mice, and nematodes, demonstrating important contributions of both *cis-*only and *trans*-only changes as well as both compensating and reinforcing joint effects of *cis*- and *trans*-acting changes (Wittkopp et al. 2004; Landry et al. 2005; McManus et al. 2010; Goncalves et al. 2012; Bell et al. 2013; Graze et al. 2014; Barrière & Ruvinsky 2014; Shen et al. 2014; Oka & Shiroishi 2014; Li & Fay 2017; Sánchez-Ramírez et al. 2021).

We sought to expand the logic of F1 hybrid inference of *cis*- and *trans*-acting divergence to investigate regulatory divergence in homozygous hybrid introgression lines of the nematodes *C. briggsae* and *C. nigoni* that show male sperm fertility defects. X-linked introgressions from *C. briggsae* into a *C. nigoni* genomic background represent genetic perturbations that may disrupt the transcriptome in ways that permit inference of *trans*-only and joint *cis-trans* regulatory divergence affecting genes in distinct ways. With this goal, we re-analysed the transcriptome dataset of Li et al. (2016) for males of two such hybrid introgression lines for which ∼95% of the genome derives from *C. nigoni*, including all autosomes. Spermatogenesis related genes are downregulated on the autosomes of these lines and 22G RNAs targeted to spermatogenesis genes are upregulated, supporting an incompatible interaction between the X-chromosome and autosomes that involves perturbation of small RNA-mediated regulation (Li et al. 2016). Here we augment these observations by inferring the gene regulatory divergence involving *cis*- and *trans*-acting changes on a per-locus basis that underlie X-autosome interactions and contribute to gene misexpression.

## RESULTS

### Independent X-linked introgressions disrupt the transcriptome in similar ways

To investigate the impacts that non-overlapping introgressions of X-linked DNA from one species (*C. briggsae*) into another (*C. nigoni*) exert on the male transcriptome of genes encoded elsewhere across the genome, we quantified differential gene expression from the data reported by Li et al. (2016). We mapped reads using reference genomes of both *C. nigoni* and *C. briggsae* to test for differential expression and misexpression in HILs relative to the parental species (cf. the analysis of Li et al. (2016), for which only the *C. briggsae* reference genome was available). We observed that most genes are expressed at similar levels across *C. nigoni, C. briggsae* and hybrid introgression lines for the shared portion of the genome, i.e., *C. nigoni* genes encoded outside of the X-linked introgression regions, consistent with prior reports (Li et al. 2016) (n_HIL1_ = 5,207 of 10,473 genes; n_HIL2_ = 4,803 of 10,541 genes). However, the HILs showed differential expression for 21% and 24% of genes in HIL1 and HIL2, respectively (n_HIL1_ = 2,229 of 10,473; n_HIL2_ = 2,543 of 10,541) (Figure 1A). The remaining ∼30% of genes in the shared genomic region either showed expression equivalent to *C. nigoni* wildtype expression (n_HIL1_ = 1124 of 10473; n_HIL2_ = 1450 of 10541) or were classified as ambiguous in each HIL (n_HIL1_ = 1913 of 10473; n_HIL2_ = 1745 of 10541). We also confirmed that both HILs showed more downregulation overall and an enrichment of downregulated genes on autosomes (Figure 1B). These downregulated genes that have lower expression in HILs than in a pure *C. nigoni* genetic background were also enriched for male specific, reproductive and germline genes (Supplementary Figure S4) (Li et al. 2016). In addition, the set of differentially expressed genes (DEGs) in HILs overlapped strongly with one another (Figure 1C, D). These similarities in differential gene expression patterns across both HILs indicate that their non-overlapping X-linked introgressions exert similar effects on the transcriptional profiles of genes encoded in the non-introgressed portion of the *C. nigoni* genome that the HILs have in common.

**Fig 1:**
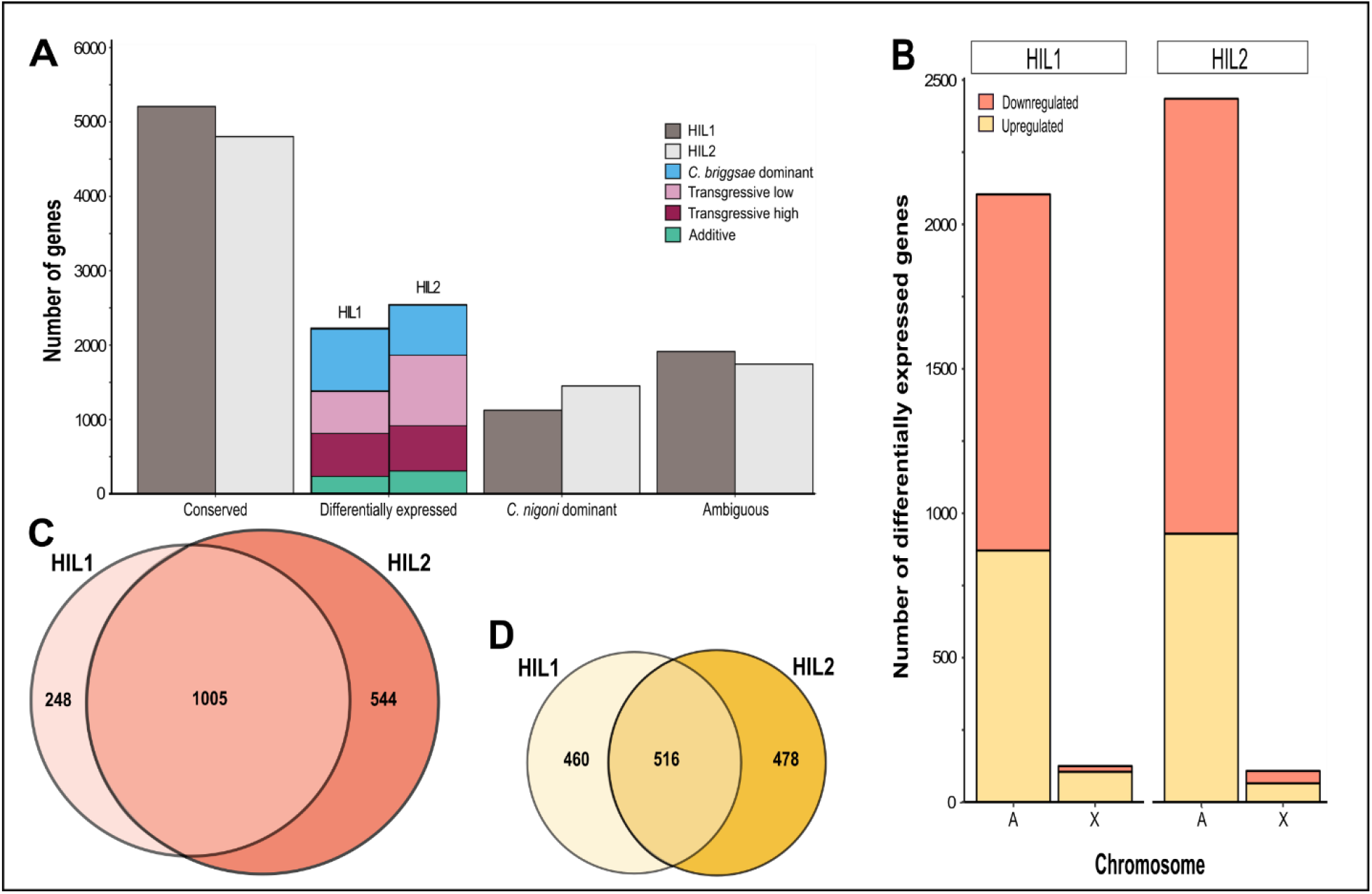
Shared patterns of differential expression by hybrid introgression lines. **(A)** Distribution of genes among differential expression designations for genes encoded within the portion of the genome shared by the HILs (i.e., excluding genes in the X-linked introgression regions). HIL2 (strain ZZY10307) included 68 more genes than HIL1 (strain ZZY10330) with detectable expression in this shared portion of the genome (n_HIL1_ = 10473; n_HIL2_ = 10541). Most genes showed a conserved expression profile in both HILs (n_HIL1_ = 5207 of 10473; n_HIL2_ = 4803 of 10541), with another 11% and 14% of genes with expression equivalent to *C. nigoni* wildtype expression in HIL1 and HIL2, respectively (n_HIL1_ = 1124 of 10473; n_HIL2_ = 1450 of 10541). Differentially expressed genes constituted 21% and 24% of total genes expressed in the common genome for HIL1 and HIL2 respectively (n_HIL1_ = 2229 of 10473; n_HIL2_ = 2543 of 10541). However, 18% and 17% of the genes in HIL1 and HIL2, respectively, did not fall into any of these categories and were classified as ambiguous (n_HIL1_ = 1913 of 10473; n_HIL2_ = 1745 of 10541). **(B)** Downregulated differentially expressed genes (orange) are more abundant than upregulated genes (yellow) in the common genomic region across the autosomes in both HILs, with few differentially expressed genes linked to non-introgressed regions of the X-chromosome (n_HIL1_ = 1253 downregulated genes of 2229 DEGs among 10473 total genes; n_HIL2_ = 1549 downregulated genes of 2543 DEGs among 10541 genes). Up- and downregulated differential gene expression is defined based on expression in HILs relative to *C. nigoni*. Downregulated genes are enriched on autosomes relative to the X-chromosome in both HILs (Fisher’s exact test P_HIL1_ < 2.2 × 10^−16^; P_HIL2_ = 7.1 × 10^−6^). **(C)** Of the 1253 and 1549 downregulated genes present on the shared genomic region in HIL1 and HIL2, respectively, a majority of genes overlapped in identity between HIL1 and HIL2 (1005 of 1253 downregulated genes of HIL1 or 1005 of 1549 downregulated genes of HIL 2; Fisher’s exact test P_overlap_ < 0.001; Jaccard Index = 0.6). **(D)** Of the 976 and 994 genes upregulated in HIL1 and HIL2, respectively, a majority overlapped in identity between HIL1 and HIL2 (516 of 976 upregulated genes in HIL1 or 516 of 994 upregulated genes in HIL2; Fisher’s exact test P_overlap_ < 0.001; Jaccard index = 0.4).

Prior work demonstrated that the X-linked introgressions in these HILs drive changes in post-transcriptional regulation through 22G smallRNAs that target spermatogenesis genes to yield greater downregulation on autosomes, ultimately resulting in hybrid male sterility (Li et al. 2016). We next sought to complement these findings by investigating the role of transcriptional regulatory divergence in hybrid male sterility, inspired by the logic of prior work on regulatory divergence inference in hybrids (Wittkopp et al. 2004; Meiklejohn et al. 2014; Mack & Nachman 2017). Specifically, we aimed to determine the relative incidence of *cis-* and *trans-*acting, as well as non-additive, regulatory divergence as revealed by perturbed patterns of gene expression caused by the non-overlapping X-linked introgressions.

### Non-overlapping introgressions produce similar patterns of transgressive expression

We first explored the additivity of inheritance of differentially expressed genes in hybrids. Based on the three-way comparison of gene expression for each HIL relative to *C. nigoni* and *C. briggsae*, we classified genes into five distinct inheritance categories: 1) *C. nigoni* “dominant” and 2) *C. briggsae* “dominant” genes showed expression levels in the HIL equivalent to one parent species, 3) “additive” genes showed expression in a HIL intermediate between both parents, and 4) “transgressive high” and 5) “transgressive low” genes showed expression in a HIL that exceeded the most extreme expression level of the parental species. Overall, we observed that both HILs showed similar profiles of expression inheritance across the genome (Figure 2A, B). For the subset of genes that were differentially expressed in both HILs across the shared portion of the genome, 84% were classified in the same inheritance category (n=1271 of 1521) (Figure 2C). In addition, over half of genes that were downregulated in HILs relative to *C. nigoni* showed transgressive-low profiles of expression (n=486 of 827 genes) and over half of upregulated genes exhibited a transgressive-high expression profile (n=257 of 444 genes). We observed qualitatively similar patterns when separately considering autosomal genes alone or X-linked genes alone that are encoded outside of the X-linked introgression regions (Supplementary Table S3). These observations indicate that 1) the X-linked hybrid introgressions lead to a prevailing signature of transgressive-low and transgressive-high expression for genes encoded elsewhere in the genome and 2) this signature affects most of the same sets of genes for both HILs.

**Fig 2:**
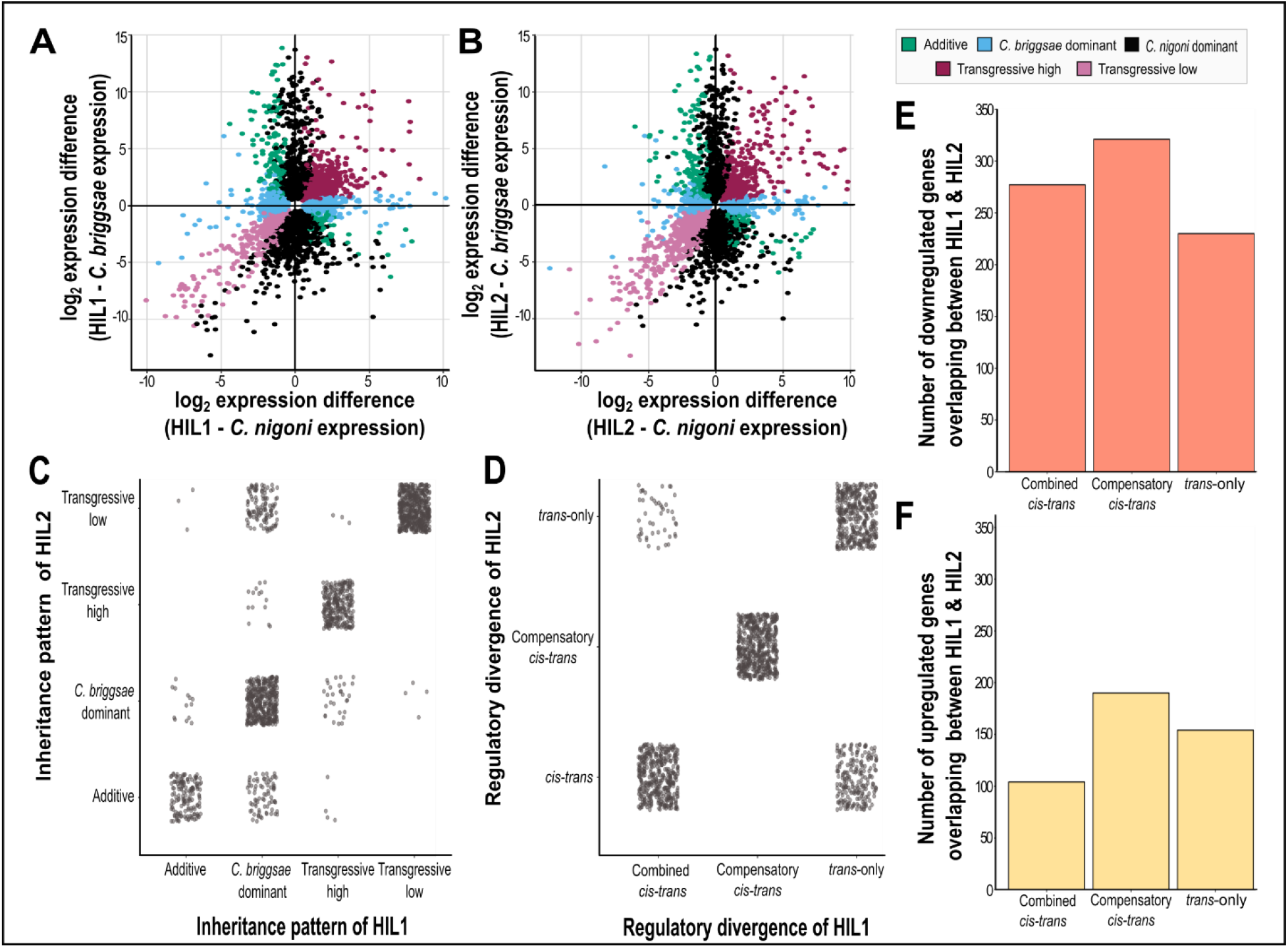
Inheritance and regulatory divergence underlying misexpression in hybrid introgression lines. **(A, B)** Scatter plots show log_2_ expression differences between each HIL and each parent species with genes coloured according to inheritance category, including only those genes in the non-introgressed portion of the genome. Genes with conserved regulation not shown for clarity. **(C)** Among the 1521 genes that were differentially expressed in both HILs, 84% corresponded to the same inheritance category in both HILs (n = 1271, points on the diagonal). Overall, 10361 genes had detectable expression in both HILs (n_HIL1_ = 10361 of 10473; n_HIL2_ = 10361 of 10541). **(D)** The regulatory divergence profiles were inferred to be equivalent for 84% of genes that were differentially expressed in both HILs (n = 1276 out of 1521 DEGs). **(E)** Out of the 1005 genes downregulated in both HILs, 59% of genes showed evidence of both *cis-* and *trans*-acting regulatory divergence in their expression profiles (n_compensatory_ = 321, n_*cis-trans*_ = 277). **(F)** Out of the 516 genes upregulated in both HILs, 67% of genes showed evidence of both *cis-* and *trans*-acting regulatory divergence in their expression profiles (n_compensatory_ = 190, n_*trans-*only_ = 154).

### Extensive overlap of cis- and trans-acting regulatory divergence revealed by distinct X-linked introgressions

We next partitioned genes into distinct categories that define alternative modes of regulatory divergence between gene orthologs of *C. nigoni* and *C. briggsae*. Inspired by the logic used to infer *cis-* and *trans-* regulatory divergence in F1 hybrids (McManus et al. 2010), we used gene expression in HILs relative to their parental species to identify loci showing 1) *trans-*only regulatory divergence, 2) compensatory *cis-* and *trans-*regulatory divergence, and 3) combined *cis-* and *trans-*regulatory divergence (which may be caused by reinforcing and/or compensatory *cis*- and *trans*-acting effects). Reminiscent of the pattern of non-additive inheritance, we observed the same regulatory divergence profiles for 84% of the differentially expressed genes that overlapped between the HILs (n = 1276 out of 1521 DEGs) (Figure 2D). Among these 1276 genes, downregulation in the HILs relative to *C. nigoni* was primarily due to divergence in both *cis-* and *trans-*acting factors (both “compensatory” and “combined”), with *trans-*only divergence providing a less frequent source of regulatory divergence (Fisher’s exact test P < 0.001, Figure 2E). In the case of upregulated genes, by contrast, misexpression primarily resulted from compensatory *cis-trans* divergence and *trans-*only regulatory divergence (Figure 2F). These observations indicate that partially-distinct regulatory controls might drive the different types of differential gene expression of hybrid introgression lines: *trans-*only divergence is more salient among upregulated genes relative to combined *cis-trans* effects, while compensatory *cis-trans* regulatory divergence contributes importantly to genes with both upregulated and downregulated expression in HILs. The extensive overlap in regulatory divergence profiles between the two HILs, however, indicates that their non-overlapping X-linked introgressions tap into a shared *trans-* acting regulatory module, providing evidence for a mutual mechanism underlying the mis-regulation and misexpression of genes in hybrids irrespective of the particular region introgressed on the X-chromosome.

## DISCUSSION

We aimed to decipher the role that gene regulatory divergence might play as a mechanism mediating X-autosome interactions that, in turn, could contribute to hybrid male sterility in *Caenorhabditis* nematodes. Using the gene expression data of two hybrid introgression lines (Li et al. 2016), we identified hundreds of genes showing distinct classes of non-additive inheritance and regulatory divergence in hybrids. High overlap across both HILs for differentially expressed genes, sets of transgressive genes, and contributions to *cis*- and *trans*-regulatory divergence together suggest that the genetically non-overlapping X-linked introgressions disrupt the transcriptome in similar ways in these hybrids, predominantly due to the joint effects of *cis*- and *trans*-acting regulatory divergence that exert compensatory effects on gene expression.

### Regulatory divergence and speciation

Conceptual arguments and empirical evidence support the idea that *cis*-acting regulatory divergence between species is common, likely facilitated by small pleiotropic effects of mutations to *cis*-regulatory elements (Stern 2000; Prud’homme et al. 2007; Wray 2007; Mack & Nachman 2017). Our analysis of homozygous hybrid introgression lines, however, is blind to detection of *cis*-only differences, letting us focus on the incidence of *trans*-only and joint *cis*- and *trans*-acting regulatory divergence. We documented how divergence in both *cis-* and *trans*-acting factors that affect a given gene’s expression provide a common source of misexpression in hybrid introgression lines, outnumbering *trans*-only effects by two-to-one. These observations contribute to growing support for a substantial contribution of the combined influence of *cis*- and *trans*-acting regulatory divergence affecting a given gene’s expression, as revealed in inter-species hybrids (Landry et al. 2005; Takahasi et al. 2011; Goncalves et al. 2012; Verta et al. 2016; Metzger et al. 2017; Mack & Nachman 2017; Sánchez-Ramírez et al. 2021). Because the regulation of gene expression involves interactions between loci, disruption of gene regulation from the joint influence of *cis*- and *trans*-acting factors can present itself as Dobzhansky-Muller expression incompatibilities if they cause fitness deficits in hybrids, and thus can be a mechanism underlying post-zygotic reproductive isolation (Mack & Nachman 2017). The finding that compensatory regulatory evolution between species is pervasive, as also demonstrated in this study, provides support for the idea that regulatory divergence can contribute to reproductive isolation as an important source of hybrid incompatibility (Meiklejohn et al. 2014; Mack & Nachman 2017; Signor & Nuzhdin 2018).

### Stabilizing selection on overall expression despite molecular evolution of gene regulation

Stabilizing selection represents the prevailing evolutionary force underlying gene expression evolution and, in particular, the pattern of widespread conservation in expression of orthologous genes between species (Gilad et al. 2006). Stabilizing selection on expression level as a phenotype can reflect different underlying mechanisms at the molecular level, however, including the selective elimination of deleterious mutations by purifying selection and/or compensatory evolution that involves distinct regulatory changes that exert counter-acting increasing and decreasing effects that jointly act to maintain overall expression levels (Mack & Nachman 2017; Signor & Nuzhdin 2018).

Our analysis provides evidence for the latter mechanism of compensatory regulatory evolution as an important molecular incarnation of stabilizing selection. We show that compensatory *cis-trans* regulatory divergence is a common feature of misexpression in hybrids, implicating stabilizing selection via compensatory regulatory divergence in the conservation of gene expression levels between species (Lemos et al. 2005; Coolon et al. 2014; Hodgins-Davis et al. 2015; Signor & Nuzhdin 2018). Such compensatory changes can arise specifically via serial changes to *cis*- and *trans-*acting regulatory factors in an evolutionary feedback within gene regulatory networks that acts to reduce the deleterious pleiotropic side effects of mutations elsewhere (e.g., modifying negatively pleiotropic side effects of a net beneficial mutation) (Maisnier-Patin & Andersson 2004; Angst & Hall 2013; Wang et al. 2015; Signor & Nuzhdin 2018). Our results add to the growing literature to support the idea that stabilizing selection operates as a major selective regime on gene expression and that it can yield interspecies expression similarity in spite of profound molecular evolution in gene regulatory controls (Landry et al. 2005; Gilad et al. 2006; Takahasi et al. 2011; Goncalves et al. 2012; Wang et al. 2015; Verta et al. 2016; Metzger et al. 2017; Sánchez-Ramírez et al. 2021). More broadly, such pervasive compensatory *cis-trans* regulatory divergence represents a key evolutionary agent for developmental system drift and as a potential source of Dobzhansky-Muller incompatibilities (True & Haag 2001; Mack & Nachman 2017; Cutter & Bundus 2020).

### Multi-locus expression incompatibilities in hybrid male sterility

The spermatogenesis process in nematode males, which is often disrupted in interspecies hybrids, involves an interconnected network of genes, with important interactions between autosomal and X-linked loci (Li et al. 2016; Bundus et al. 2018). Our results, and those of Li et al (2016), show that distinct introgression segments on the X-chromosome disrupt in similar ways the transcriptome of autosomally-encoded genes. By affecting the same spermatogenesis-related genes on the autosomes, a broad region of the X-chromosome is capable of perturbing proper spermatogenesis to cause infertility (Li et al. 2016). This overlapping sensitivity of the autosomal transcriptome to different X-linked genetic perturbations implicates the presence of multi-way negative genetic interactions as a complex form of Dobzhansky-Muller expression incompatibility.

Evolution in partially-redundant gene regulatory networks (GRN) can lead to compensatory molecular evolutionary changes that nonetheless retain a constant phenotype, termed developmental system drift (True & Haag 2001; Schiffman & Ralph 2022). Consequently, as different mutations accumulate independently in separate species, developmental system drift can lead their molecular genetic pathways to trace distinct trajectories of connectivity, despite reaching the same phenotype. In hybrids, however, the crosstalk between divergent pathways can lead to genetic mismatches. Such mismatches can disrupt epistatic interactions between the coevolved expression of loci in the GRN, a form of Dobzhanky-Muller incompatibility that may lead to hybrid dysfunction (Tulchinsky et al. 2014; Satokangas et al. 2020). Because *cis-* and *trans-*acting regulatory controls within a GRN jointly serve to mediate multi-locus epistatic interactions through pairwise or higher order complexes, such regulatory divergence can influence the likelihood that genetic incompatibilities arise between species and contribute to reproductive isolation (Satokangas et al. 2020).

Simple pairwise interactions that involve a given locus, however, should not be influenced directly by multiple distinct genetic perturbations. We found that similar disruptions to the transcriptome emerge in hybrid introgression lines due to distinct, non-overlapping genetic perturbations of the X-chromosome, often mediated by compensatory *cis*- and *trans*-acting regulatory divergence. Consequently, these observations require the existence of multi-way incompatibility interactions that involve at least three, but perhaps many more, genes (Burkart-Waco et al. 2012). Thus, the GRN responsible for male fertility in this system may be structured in a way that makes it susceptible to disruption in hybrids as a result of multi-way incompatibilities. Given that the evolutionary dynamics of multi-way DMIs differ from pairwise DMIs (Orr 1995; Kondrashov 2003), it remains an important outstanding question to determine whether regulatory divergence in GRNs is predisposed more generally to producing multi-way DMIs. Such differences among the GRNs that control the development of different traits might help explain why some aspects of development are more prone to manifesting DMIs and hybrid dysfunction (Cutter & Bundus 2020). If GRNs that control the developmental programs responsible for male fertility are disproportionately prone to multi-way DMIs, then such genetic architectures might provide a mechanistic rationale for the “fragile male” hypothesis to explain Haldane’s rule (Wu & Davis 1993; Cutter 2018).

## METHODS

### Dataset

We accessed the mRNA sequence dataset produced by Li et al. (2016) from NCBI (SRP067756). This dataset consists of ∼8 million high quality paired-end (2×150bp) Illumina MiSeq reads per sample from pools of 300 young adult males, with replication in triplicate for each of *C. nigoni* (JU1421), *C. briggsae* (AF16), ZZY10330 (hybrid introgression line 1), and ZZY10307 (hybrid introgression line 2) (Li et al. 2016). The genomes of hybrid introgression lines (HILs) primarily carry *C. nigoni* DNA, with each HIL containing an independent, non-overlapping fragment from the *C. briggsae* X-chromosome that contributes to hybrid male sterility (Bi et al. 2015; Li et al. 2016). In *Caenorhabditis* males, the X-chromosome is hemizygous due to the XO sex-determination system. Based on our analysis of mapped reads below, we affirmed that the right arm of the X-chromosome in ZZY10330 (HIL1) carries a ∼4.8Mb fragment from *C. briggsae* containing a total of 311 orthologous genes with detectable expression inside the fragment (out of total 470 annotated orthologous genes) and 11,097 autosomal and X-linked orthologous genes with detectable expression outside the introgression from the *C. nigoni* genomic background (from a total 13,505 annotated orthologous genes outside the introgressed region). The ZZY10307 (HIL2) strain has an introgression fragment of size ∼7.4 Mb in the middle of the X-chromosome with a total of 631 orthologous genes inside the introgression (out of total 867 annotated orthologous genes) and 10,852 outside the introgression with detectable expression.

### Analysis of mRNA-seq reads

We mapped the same mRNA reads separately against both the *C. nigoni* (JU1422) reference genome “nigoni .pc_2016.07.14” (Yin et al. 2018) and the *C. briggsae* (AF16) reference genome “CB4” (Ross et al. 2011) using STAR with default parameters (Dobin et al. 2013). To identify and confirm the introgression boundaries on the X-chromosome, for each HIL, we compared read counts mapped to *C. briggsae* and *C. nigoni* reference genomes to determine which genes mapped better to the *C. briggsae* genome as a consequence of ∼21% synonymous-site sequence divergence of *C. nigoni* orthologous loci (Thomas et al. 2015). For each HIL, we calculated and plotted the difference in log_2_(mean read count + 0.1) when mapped to the *C. briggsae* reference genome versus the *C. nigoni* reference genome for each gene on the X-chromosome (Supplementary Figure 1, 2; Supplementary Table S1, S2). Genes with positive values for this difference indicated better mapping to the *C. briggsae* reference genome, clustering at the regions expected to contain the introgression which we then used to define the introgression boundaries for each HIL. In this way, we confirmed the presence of a ∼4.8Mb fragment introgressed on the right arm of the X-chromosome in HIL1 from *C. briggsae* (*C. briggsae* positions 16,392,964 bp to 21,277,039 bp, defined by the genes WBGene00041261 on the left and WBGene00031970 on the right of the fragment) (Supplementary Table S1) and a ∼7.4 Mb fragment in the middle of the X-chromosome of HIL2 (*C. briggsae* positions 4,742,872 bp to 12,165,712 bp, defined by the genes WBGene00036950 on the left and WBGene00032130 on the right of the introgressed fragment) (Supplementary Table S2).

To quantify gene expression appropriately for genes inside and outside the introgression region, we considered mapped reads to the *C. nigoni* reference genome for those genes determined to be outside the introgression region of a given HIL, and mapped reads to the *C. briggsae* reference genome for those genes inside each introgression. We quantified gene expression of all the genes using FeatureCounts (Liao et al. 2014) and restricted subsequent analysis of the read count data to overlap with the set of 13,975 1:1 orthologous genes defined in Sánchez-Ramírez et al. (2021). Differential gene expression analysis was performed for orthologous genes using DESeq2 (Love et al. 2014) in R Studio (RStudio version 4.1.2). Genes showing Benjamini-Hochberg false discovery rate-adjusted P_adj_ < 0.05 between *C. nigoni* and a given HIL were considered to be differentially expressed. We then categorized genes from their expression profiles to infer additivity of inheritance pattern and *cis-trans* regulatory divergence, similar to McManus et al. (2010). Genes were classified into five distinct inheritance categories based on their expression relative to the wildtype expression of both *C. briggsae* and *C. nigoni*: 1) and 2) Genes showing expression levels in the HIL equivalent to one of the parent species were classified as *C. nigoni* (or *C. briggsae)* “dominant”, 3) genes showing expression in a HIL significantly different from and intermediate between both parents were termed “additive”, 4) and 5) genes showing expression in a HIL that was significantly above or below both parental species were classified as genes with “transgressive high” or “transgressive low” expression, respectively (Supplementary Figure S3B).

To infer distinct kinds of regulatory divergence, we classified genes according to distinct profiles of differential expression in pairwise contrasts between each of *C. briggsae, C. nigoni*, and hybrid introgression lines. In particular, we inferred genes to exhibit 1) *trans-*only regulatory divergence when *C. nigoni* expression differed significantly from *C. briggsae* and the HIL in the absence of significant differential expression of the HIL from *C. briggsae*, 2) compensatory *cis-trans* regulatory divergence when the HIL showed significant differential expression with both *C. nigoni* and *C. briggsae* despite *C. nigoni* and *C. briggsae* displaying no significant difference between one another, 3) combined *cis-* and *trans-* regulatory divergence when all three pairwise contrasts displayed significant expression differences (because this category includes genes in HILs showing expression higher or lower than both parents as well as genes showing intermediate expression between parents, the *cis-trans* effects could be either compensatory or reinforcing), 4) conserved regulation when the gene showed no significant differential expression in any pairwise contrast, or 5) ambiguous regulatory divergence for all other expression circumstances (Supplementary Figure S3C). It is important to note that although our logic for classification is based on McManus et al. (2010), and Meiklejohn et al. (2014) that exploited heterozygous autosomal introgressions in *Drosophila*, our criteria differ to account for the genomic architecture of hemizygous X-linked introgressions of hybrid males in this study, rather than allele-specific expression of F1 hybrids.

## Acknowledgements

This work was supported by the Natural Sciences and Engineering Research Council of Canada with a Discovery Grant to ADC. We thank Santiago Sánchez-Ramírez for help with the differential gene expression analysis and inference of regulatory divergence as well as Katja Kasimatis, Rebecca Schalkowski, and Daniel Fusca for helpful discussion and comments on the manuscript.

## Data Availability

### No new data were generated in support of this work

The mRNA sequencing data underlying this article are available in NCBI Gene Expression Omnibus at GEO; http://www.ncbi.nlm.nih.gov/geo/ and can be accessed with accession number GSE76306. The *C. nigoni* and *C. briggsae* reference genomes used in this study can be accessed under the NCBI BioProject accessions PRJNA384657 and PRJNA10731 respectively.

