## Supplementary Figures and Captions for "Regulatory divergence as a mechanism for X-autosome incompatibilities in *Caenorhabditis* nematodes"

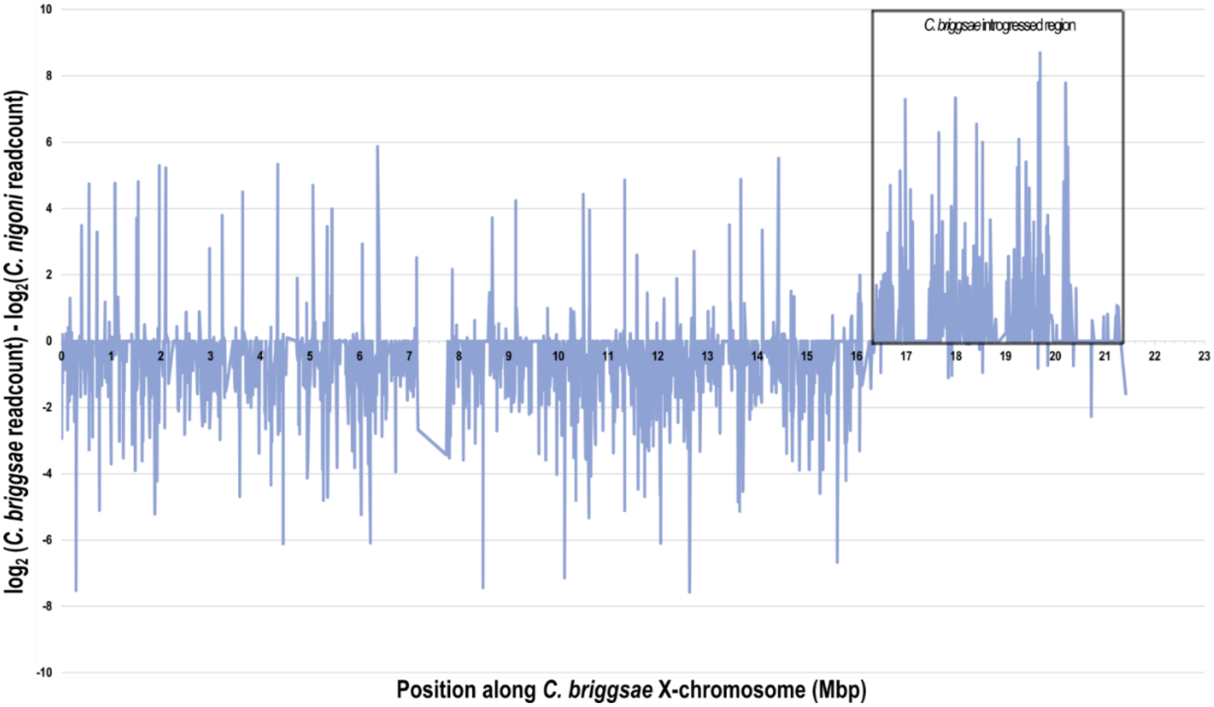


**Supplementary Figure S1: Confirmation of introgression boundary for HIL1 (strain ZZY10330).** Plot showing difference in $\log_2$(mean read count + 0.1) between readcounts obtained from mapping using *C. briggsae* reference genome and readcounts obtained from mapping using the *C. nigoni* reference genome for each gene along the X-chromosome (*C. briggsae* position). A positive value for this difference between readcounts indicates that the gene maps better to *C. briggsae* reference genome and hence is likely part of the C. *briggsae* X-chromosome i.e., part of introgressed region. A cluster of positive values around the expected introgression region (region within black box) was observed on the right arm of the X-chromosome. This region was used to define the introgression boundaries and confirmed the presence of ~4.8Mb fragment (*C. briggsae* positions 16.39 Mb to 21.26 Mb) defined by the genes WBGene00041261 at 16392964bp on the left and WBGene00031970 at 21269250bp on the right of the fragment introgressed in HIL1 from *C. briggsae*.

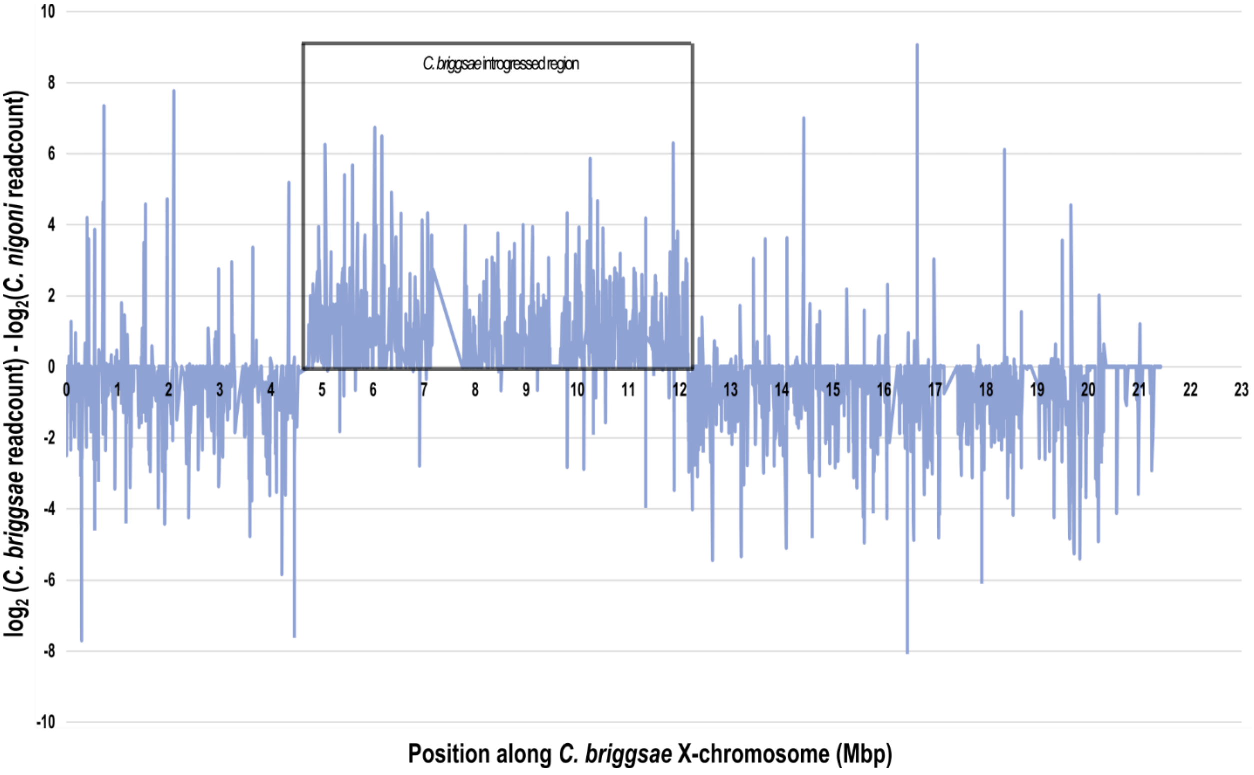


**Supplementary Figure S2: Mapping introgression boundary for HIL2 (strain ZZY10307).** A cluster of positive values was observed in the middle of the X-chromosome (region within black box). This region confirmed the presence of ~7.4Mb fragment (*C. briggsae* positions 4.74 Mb to 12.16 Mb) defined by the genes WBGene00036950 at 4742872bp on the left and WBGene00032130 at 12160549bp on the right of the *C. briggsae* fragment introgressed in HIL2.

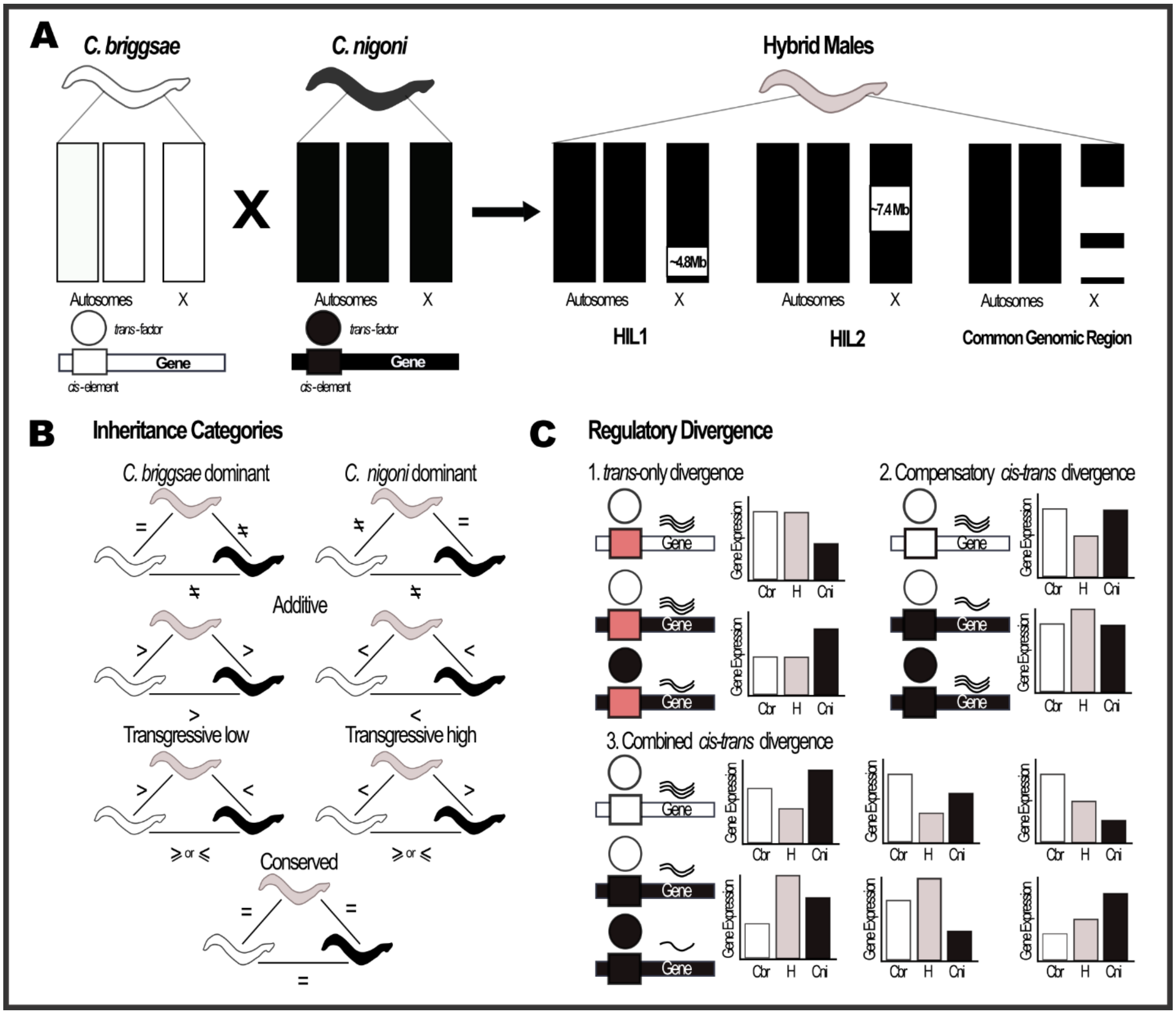


**Supplementary Figure S3: Criteria for classification of inheritance and regulatory divergence categories for genes in the shared genomic region. (A)** HIL1 and HIL2, respectively, have ~4.8Mb and ~7.4Mb regions on the X-chromosome introgressed from *C. briggsae* (white) in *C. nigoni* background (black). Different *cis*-elements (rectangles) and *trans*-acting factors (circles) may be encoded by *C. briggsae* (white) and *C. nigoni* (black). **(B)** Using differential gene expression analysis, genes were classified into five different inheritance categories based on a three-way comparison of expression between *C. briggsae*, *C. nigoni* and hybrids: 1) Genes showing expression equal to one of the wildtype species that also showed significant difference between the two parent species were termed *C. briggsae* or *C. nigoni* “dominant”, 2) Genes showing expression significantly different between the parents and significantly different but intermediate between both parents in HIL were termed “additive”, 3) and 4) genes showing expression significantly different and below or above both wildtype species were termed “transgressive low” and “transgressive high,” respectively, and 5) genes showing no significant differences in expression relative to wildtype gene expression were termed conserved. All other expression patterns were termed “ambiguous”. Equal signs indicate non-significant expression differences. **(C)** Pairwise contrasts of differential gene expression analysis between *C. briggsae*, *C. nigoni*, and HILs were used to infer three distinct regulatory divergence profiles. 1) *C. nigoni* expression differing significantly from *C. briggsae* and the HIL, in the absence of significant expression difference between HIL and *C. briggsae* were classified as *trans*-only regulatory divergence. This is because when genes in *C. nigoni*, *C. briggsae*, and the HIL differ only in the *trans*-acting factor and have a common *cis*-element (pink), the underlying *cis*-*trans* interaction in HILs is similar to interaction within *C. briggsae*. This category may underestimate the incidence of joint *cis*-*trans* regulatory divergence because, in some cases, there could exist *cis*-regulatory divergence that could contribute to this pattern. 2) When the gene in the HIL showed significant differential expression with both *C. nigoni* and *C. briggsae* genes, despite *C. nigoni* and *C. briggsae* displaying no significant difference between one another, it was classified as compensatory *cis*-*trans* regulatory divergence. 3) When all three pairwise contrasts displayed significant expression differences from one another, it was classified as combined *cis*- and *trans*-regulatory divergence. These combined effects could be compensatory, reinforcing, or a mixture of both. When the gene showed no significant differential expression in any pairwise contrast its regulation was classified as conserved and all other expression circumstances were classified as ambiguous regulatory divergence.

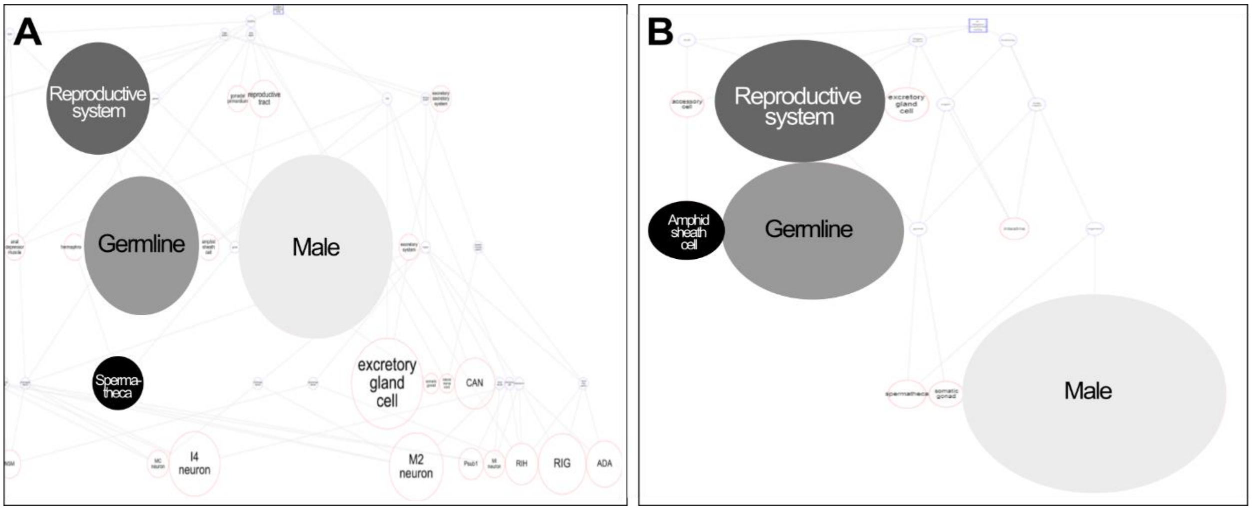


**Supplementary Figure S4: Tissue enrichment analysis (TEA) of downregulated genes in the shared genomic region**. The downregulated genes that have lower expression in HILs than in a pure *C. nigoni* genetic background were also enriched for male specific, reproductive and germline genes. Tissue enrichment analysis of downregulated genes in HIL1 (strain ZZY10330) (A) and HIL2 (strain ZZY10307) (B) reveal patterns that are consistent with previous analysis by (Li et al. 2016). TEA conducted using Wormbase enrichment analysis ($n_{HIL1}$ = 1253 downregulated genes of 2229 DEGs among 10473 total genes; $n_{HIL2}$ = 1549 downregulated genes of 2543 DEGs among 10541 genes) (Angeles-Albores et al. 2016, 2018).

### Supplementary Tables

All described tables are available in the file Viswanath-Cutter_Supplementary-Tables_24-5-2022.xlsx.

### Supplementary Table legends

**Table S1: Readcounts used to identify introgression boundaries of HIL1 strain ZZY10330**

Table contains readcounts for X-linked genes in all three replicates of *C. briggsae*, *C. nigoni,* and HIL1. For genes in HIL1, readcounts were obtained by mapping using both *C. briggsae* and *C. nigoni* reference genomes. The difference in mapped reads between them ($log_2$(mean readcount + 0.1)) was used as a metric to identify introgression boundaries, with positive values indicating that the gene mapped better to *C. briggsae* than *C. nigoni* and hence is likely part of the introgressed region. The genes marking the boundary of the introgressed region are highlighted

**Table S2: Readcounts used to identify introgression boundaries of HIL2 strain ZZY10307**

Table contains readcounts for X-linked genes in all three replicates of *C. briggsae*, *C. nigoni,* and HIL2. For genes in HIL2, readcounts were obtained by mapping using both *C. briggsae* and *C. nigoni* reference genomes. The difference in mapped reads between them ($log_2$(mean readcount + 0.1)) was used as a metric to identify introgression boundaries, with positive values indicating that the gene mapped better to *C. briggsae* than *C. nigoni* and hence is likely part of the introgressed region.

**Table S3: Autosomal and X-linked genes in the shared genomic region show similar patterns of overlap in inheritance across HILs**

Table shows the inheritance pattern across autosomal and X-linked genes present in the common genomic region across HILs. Overall, 10361 genes had detectable expression in both HILs in the shared genomic region ($n_{HIL1}$ = 10361 of 10473; $n_{HIL2}$ = 10361 of 10541) out of which 9598 genes were present on the autosomes and 763 genes were present on the X-chromosome. Overall, ~65% and ~72% of the autosomal and X-linked genes, respectively, were categorised in the same inheritance category across both HILs ($n_{autosomal\ overlap}$ = 6274 of 9598; $n_{X\text{-}linked\ overlap}$ = 551 of 763).
